# Benchmarking small variant detection with ONT reveals high performance in challenging regions

**DOI:** 10.1101/2020.10.22.350009

**Authors:** Peter L. Møller, Guillaume Holley, Doruk Beyter, Mette Nyegaard, Bjarni V. Halldórsson

## Abstract

**Background:** The development of long read sequencing (LRS) has led to greater access to the human genome. LRS produces long read lengths at the cost of high error rates and has shown to be more useful in calling structural variants than short read sequencing (SRS) data. In this paper we evaluate how to use LRS data from Oxford Nanopore Technologies (ONT) to call small variants in regions in- and outside the reach of SRS.

**Results:** Calling single nucleotide polymorphisms (SNPs) with ONT data has comparable accuracy to Illumina when evaluating against the Genome in a Bottle truth set v4.2. In the major histocompatibility complex (MHC) and regions where mapping short reads is difficult, the F-measure of ONT calls exceeds those of short reads by 2-4% when sequence coverage is 20X or greater.

We develop recommendations for how to perform small variant calling with LRS data and improve current approaches to the difficult regions by re-genotyping variants to increase the F-measure from 97.24% to 98.78%. Furthermore, we show how LRS can call variants in genomic regions inaccessible to SRS, including medically relevant genes such as *STRC* and *CFC1B*.

**Conclusions:** Although small variant calling in LRS data is still immature, current methods are clearly useful in difficult and inaccessible regions of the genome, enabling variant calling in medically relevant genes not accessible to SRS.

## Introduction

The field of genomics is constantly evolving as developments in sequencing technology allow greater access to genomic variation. Since the turn of the century, short read sequencing (SRS) has led to tremendous insight into the human genome, with SRS becoming an integral part of diagnostics [1]. Currently, SRS is almost synonymous with Illumina sequencing, with read lengths around 150 bp and error rates from 0.1-1% depending on platform and protocol [2].

In the past decade we have seen the emergence of long read sequencing (LRS), with Pacific Biosciences’ (PacBio) single-molecule real-time (SMRT) technology in 2011 and Oxford Nanopore Technologies (ONT) in 2014 [3]. Both technologies were initially plagued by high error rates (10-15%) [4], making variant calling very challenging. PacBio solved this issue with the introduction of the circular consensus sequencing (CCS) protocol, producing high fidelity (HiFi) reads with lengths of 10-20 kb and error rates of 0.2% [5]. The drawback of this approach is a highly reduced output, with a single SMRT Cell 8M producing 15-25 Gb of HiFi data [6]. Meanwhile, ONT is still error prone but provides far longer reads, typically 10-100 kb, and outputting 50-100 Gb of data per PromethION flow cell [6]. ONT also offers an ultra-long read protocol consistently producing reads exceeding 100 kb at the cost of decreased output [7]. Considering the PromethION flow cell being slightly cheaper than the SMRT Cell, this results in a much lower cost per base for ONT data [8].

The development of LRS has made previously inaccessible regions of the genome available for study [9]. These regions were described by Ebbert et al. as “dark”, due to low coverage (≤ 5 reads) or low mapping quality (≥ 90% reads with MAPQ < 10). They found that PacBio reduced the percentage of dark bases by 58.2% for all gene bodies and 77.7% for coding sequence (CDS). In comparison ONT reduced the percentage of dark bases by 77.9% in all gene bodies and by 95.6% in the CDS. Recently, the Telomere-to-Telomere (T2T) consortium also showed the strength of ultra-long ONT reads, using them as part of their effort to create a complete assembly of the human CHM13hTERT cell line, underlining the importance of read length in accessing dark regions. [10,11]. Further proof of the usefulness of LRS was the genome in a bottle (GIAB) truth set v4.1, which expanded the high confidence regions to 92.2% of the genome compared to 85.4% in v3.3.2 using PacBio HiFi reads [12].

Here we chose to evaluate variant discovery across the genome based on ONT data, as the technology provides the greatest access to the dark regions at the lowest cost per base. We test some of the most recent variant callers, namely Medaka, a diploid-aware neural network developed by ONT [13]; Clair, a deep neural network based variant caller [14] and P.E.P.P.E.R./DeepVariant (“PEPPER” going forward), a deep neural network polisher and caller [15,16] presented in the PrecisionFDA Truth Challenge V2.

## Results

### Truth set benchmarking reveals inconsistent performance across different evaluation regions

We analyzed data from the publicly available Ashkenazim trio (HG002, HG003, HG004). Variant calls were evaluated against the GIAB v4.2 truth set released in relation to the PrecisionFDA Truth Challenge V2 capturing 92.2% of the genome. Illumina data was benchmarked with DeepVariant [16] to establish baseline performance of a known caller with SRS data. The HG002 truth set has been widely used for model training, we therefore use the HG003 and HG004 truth sets for evaluation and report their average.

The Illumina data had 60X coverage for all individuals, while coverage for ONT data varied from 50X for HG002 (8.81% error rate) to 80X for HG003 (7.82% error rate) and HG004 (8.24% error rate).

As seen in Figure 1, the best variant calling performance across all benchmark regions is achieved using Illumina data. This is no surprise, as short reads were the foundation of previous versions of the truth set and most of the human genome is sufficiently unique to map short reads unambiguously. When stratifying performance by variant type, we see a more detailed picture. This highlights decent SNP calling with ONT data, with both Medaka and PEPPER F-measure surpassing 99%, while Clair achieves 98.55%. Meanwhile, indel detection with ONT lags severely behind. For Illumina we see an F-measure of 99.58%, while Medaka, the best performing ONT caller, achieves 70.34%.

**Figure 1.**
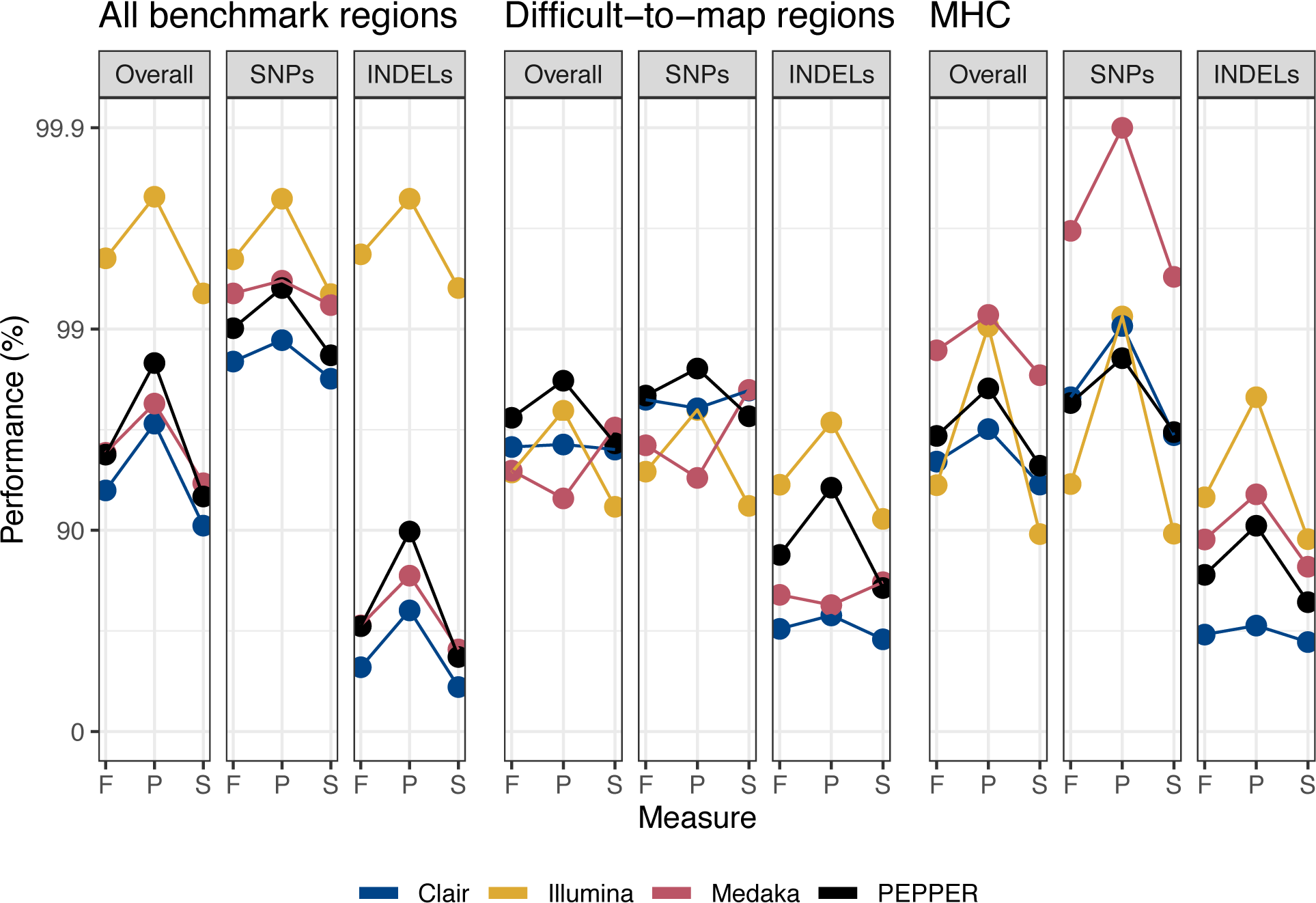
Performance metrics measured against the Genome in a Bottle v4.2 truth set. All benchmark regions: Complete truth set. MHC: Intersect of major histocompatibility complex and truth set. Difficult-to-map regions: Intersect of the truth set with segmental duplications and regions where 100 bp read pairs have <= 2 mismatches and <= 1 indel difference from another part of the genome. MHC: Major histocompatibility complex; F: F-measure; P: Precision; S: Sensitivity

Analyzing performance in more complex regions, such as the difficult-to-map regions and the major histocompatibility complex (MHC), reveals the benefit of LRS. In the difficult-to-map regions (145 mb), overall ONT performance surpasses Illumina with an F-measure of 97.24% using PEPPER, while Illumina reaches 94.84%. A similar picture is seen in the MHC (4.6 mb), except the overall best performance is achieved by Medaka, having an F-measure of 98.73%, while Illumina reaches 94.04%. In both the MHC and the difficult-to-map regions Illumina is 5-8% better for calling indels than the best ONT caller.

### Subsampling reveals high performance from 30X coverage

As 80X ONT whole genome sequencing is not necessarily feasible for large scale experiments we benchmarked performance at 10X coverage increments (Figure 2). This highlighted the need for at least 20X ONT to surpass SRS performance in the difficult-to-map regions, while 30X was necessary in the MHC to achieve a meaningful improvement. Interestingly, this also showed PEPPER as the better choice for ONT data across most depths and regions, despite the author recommendation of 50-80X coverage [15].

**Figure 2.**
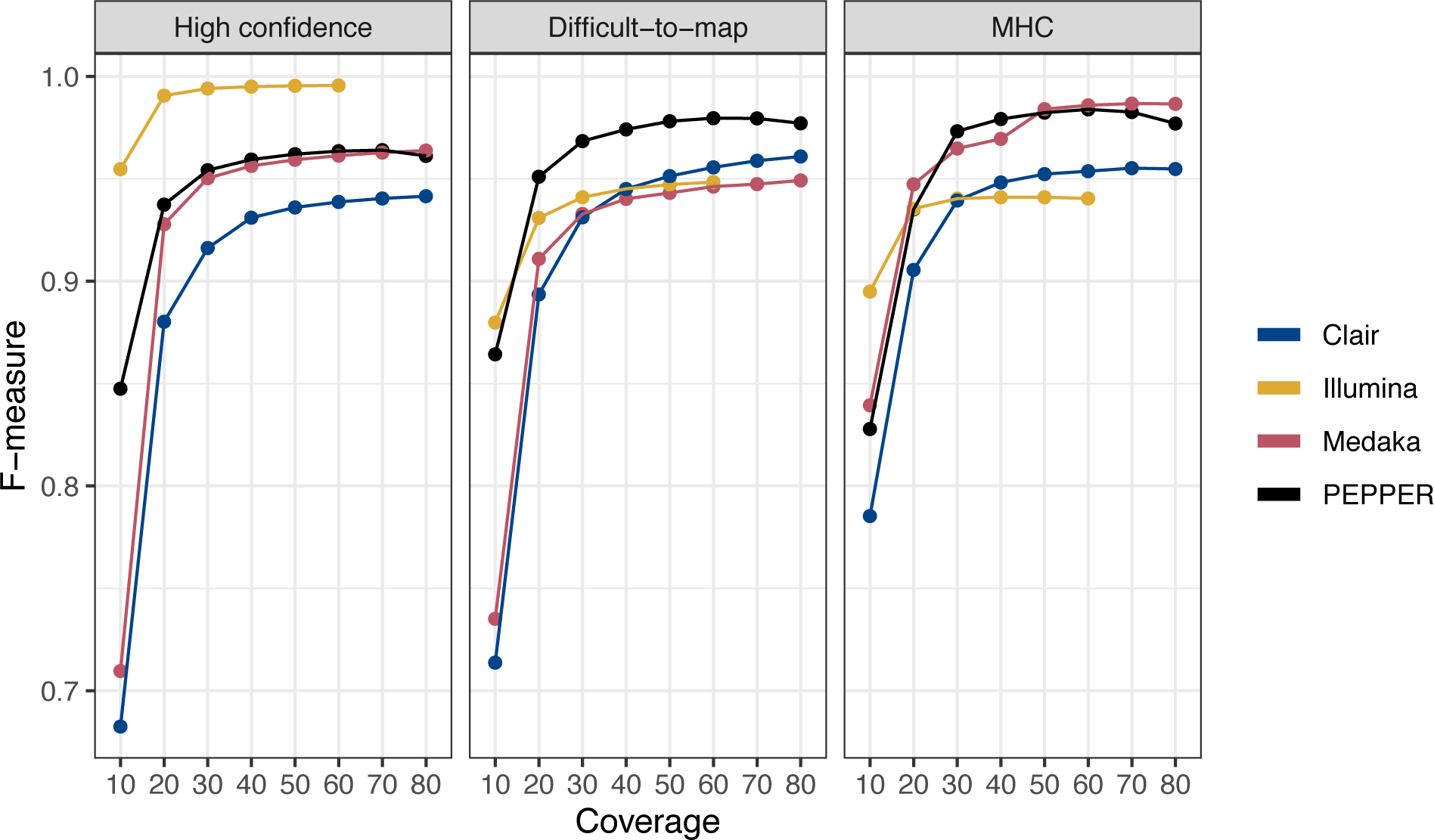
Variant calling performance as a function of sequence coverage. F-measure was determined between 10 and 80X coverage in 10X increments for each evaluation region. Numbers are average of HG003 and HG004. MHC: Major histocompatibility complex.

PEPPER with 30X coverage resulted in an overall f-measure of 95.42%, while doubling the coverage increased performance less than 1%. Finally, we observed a slight dip in PEPPER performance above 60X coverage.

### Mendelian concordance decreases outside high confidence regions

Evaluating the performance of variant callers in the 7.8% of the genome not included in the GIAB truth set is more difficult. Here we look at two measurements; 1) the total number of variants as an indicator of sensitivity and 2) the Mendelian concordance as an indicator of precision. Mendelian concordance is a commonly used metric, when no truth set exists [17], while the number of variants is important to avoid overestimating the performance of conservative callers. To achieve consistent Mendelian concordance calculations for each setup, we only benchmark variant callers supporting gVCF output (DeepVariant for Illumina, PEPPER for ONT).

As seen in Table 1, variant calling in the high confidence (HC) regions is very consistent, with both technologies resulting in Mendelian concordance above 99%. At the same time, the total number of variants in ONT data displays a shortcoming of Mendelian concordance. The call set is missing approximately 200,000 indels, caused by low sensitivity (57.41%, Figure 1), but maintains high concordance due to consistently missing indels. In the complementary (Comp) regions the mendelian concordance decreases for both technologies. Stratifying by variant type shows the concordance of ONT SNPs to be within 2% of Illumina, while calling 90,000 more SNPs, presumably due to greater access to the traditionally dark regions of the genome.

**Table 1.** Total number of variants called in HG002 and their Mendelian concordance.

| Data | Region (variant type) | Variants | MC (%) |
| --- | --- | --- | --- |
| **Illumina** | HC (SNPs + indels) | 3,872,611 | 99.87 |
|  | Comp (SNPs + indels) | 734,056 | 95.00 |
|  | Comp (SNPs) | 382,568 | 94.59 |
|  | Comp (indels) | 335,519 | 95.81 |
| **ONT** | HC (SNPs + indels) | 3,662,447 | 99.40 |
|  | Comp (SNPs + indels) | 584,514 | 91.24 |
|  | Comp (SNPs) | 469,811 | 92.62 |
|  | Comp (indels) | 109,753 | 85.62 |

*MC: Mendelian concordance; HC: High confidence; Comp: Complementary regions; SNPs: Single nucleotide polymorphisms; ONT: Oxford Nanopore Technologies.*

### Long reads reveal 22 mb of dark genome including medically relevant genes

We identified dark regions for both short and long reads, subsequently identifying regions uniquely dark to either technology. Here we adapted the dark region definition from [9] described previously. This approach found 22 mb of the genome dark only to short reads, while, surprisingly, 1.5 mb was solely dark to long reads (Figure 3). These regions were spread across 32607 sites, ranging from 1 to 103,863 bp (median 155 bp) in size, with the 1324 largest regions (3,335 bp and above) making up 50% of the dark bases.

**Figure 3.**
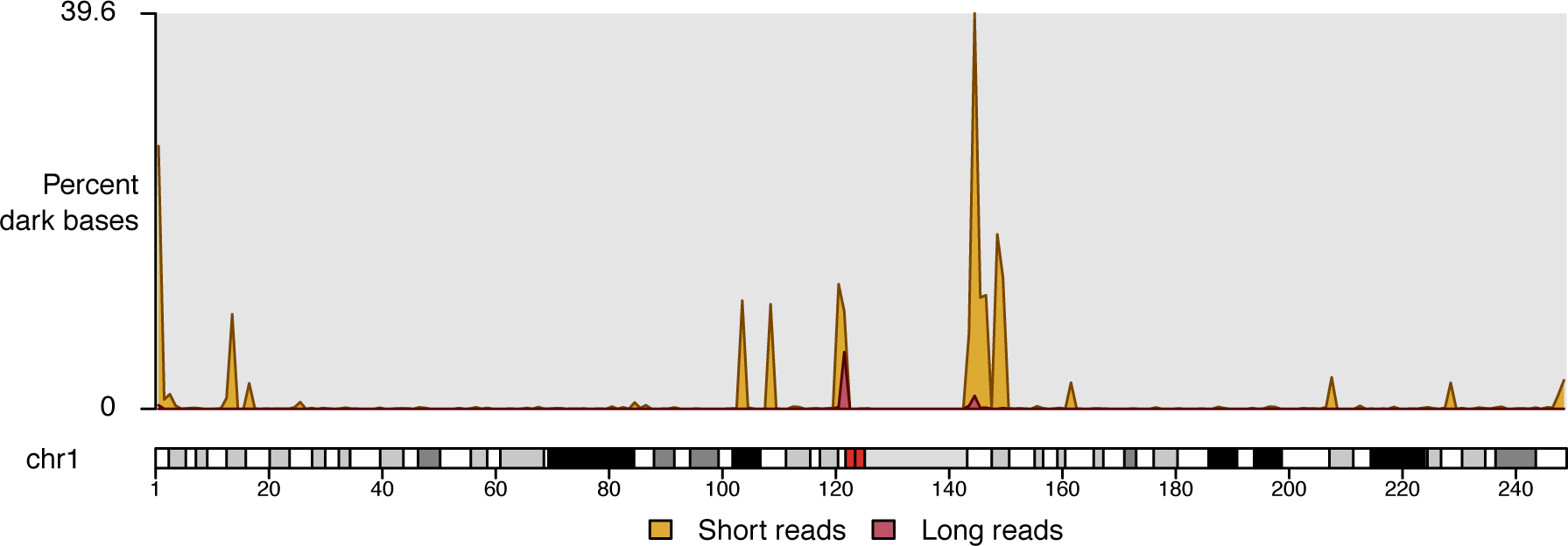
Dark regions of chromosome 1. Percentage of bases in 1 mb windows which are dark to only one technology.

Looking at genes previously defined as medically relevant but challenging to short-read technologies [18], we find 618 of 4,773 genes where some bases can only be reached by ONT and 49 genes where at least 10% of the gene is only reachable with ONT. The same approach for all genes in the Ensembl database [19] identified 2,336 of 19,190 genes with any bases only reached by ONT and 453 genes above the 10% threshold. Figure 4 shows two medically relevant genes, *STRC* and *CFC1B*, of which 23.39% and 100% of the genes can only be accessed with ONT. A comparison to PacBio HiFi found 12 mb of the genome, including *CFC1B*, to be dark to PacBio Hifi data but not ONT.

**Figure 4.**
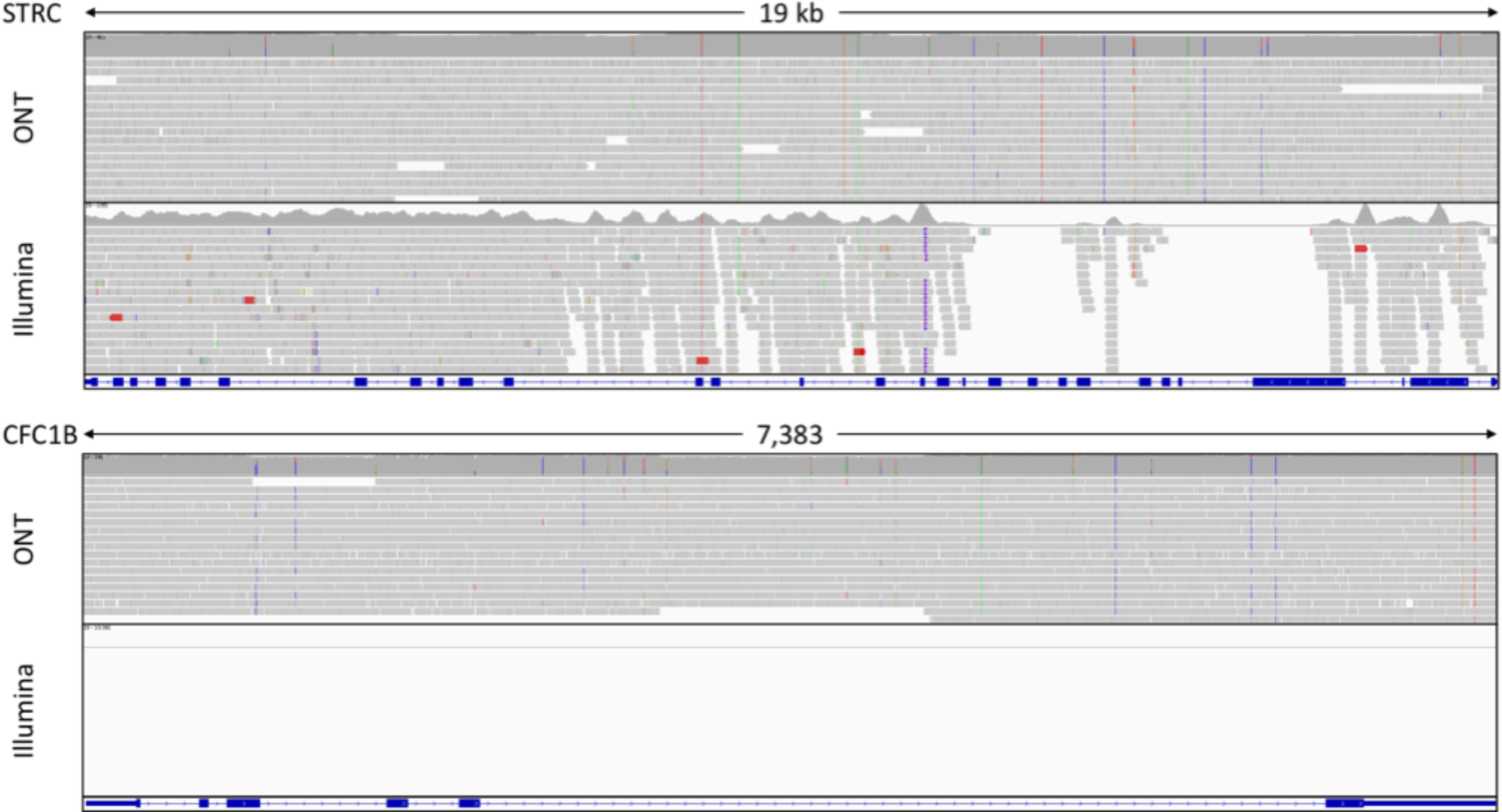
Coverage of the medically relevant genes STRC and CFC1B. Top panel shows the coverage of the STRC gene. Bottom panel shows the coverage of the CFC1B gene.

Intersecting the short read dark regions with the PEPPER variants identified 54,000 variants in HG002, of which 48,000 were SNPs. This number corresponds to half of the 90,000 SNPs identified by ONT and not Illumina. Analysis of these variants will however be difficult as Mendelian concordance is only 86.87%. Furthermore, the PEPPER call set contained an extra 25,000 events in these regions, which were not called as variants. Upon visual inspection several events were missing genotypes (./.) or called as homozygous reference (0/0) despite high coverage and variant allele frequencies above 50%. We assume this is caused by DeepVariant (the final step of PEPPER) sometimes recognizing SNP-dense regions as segmental duplications, leading it to call homozygous reference [20].

### Re-genotyping variants outside the high confidence regions increase mendelian concordance

As our original calls in the dark regions showed low Mendelian consistency, we re-genotyped the joint VCF files from PEPPER using Whatshap [21]. Whatshap can take an input VCF + BAM file to compute haplotype-aware genotypes for each event in the VCF file based on a Hidden Markov Model [22]. Using this approach on all events in the dark regions increased the number of variants by approximately 15,000 while simultaneously improving the Mendelian concordance by more than 7% (Table 2).

**Table 2.** Number of variants and Mendelian concordance of PEPPER and re-genotyped PEPPER in the dark regions.

| Variants | MC (%) | Re-genotyping |
| --- | --- | --- |
| 54,363 | 86.87 | No |
| 71,866 | 94.32 | Yes |

MC: Mendelian concordance

Extending the re-genotyping to each subset of the genome (HC, MHC, difficult-to-map, complementary, dark regions) showed increased Mendelian concordance for all regions (Figure 5). For the high confidence, MHC and difficult-to-map regions we also re-calculated the F-measure of the new HG003 and HG004 genotypes. Re-genotyping resulted in 1.5% decreased F-measure in the high confidence regions. However, in the MHC, the F-measure increased marginally, while in the difficult-to-map regions it improved from 97.25% to 98.78%. Further, re-genotyping improved the consistency of variant calling between samples in the MHC. In this region, the F-measure for both Illumina and ONT varied 2-4% between HG003 and HG004, while in other regions it varied 0-0.2%. Re-genotyping reduced the F-measure difference in the MHC to 0.2-2%.

**Figure 5.**
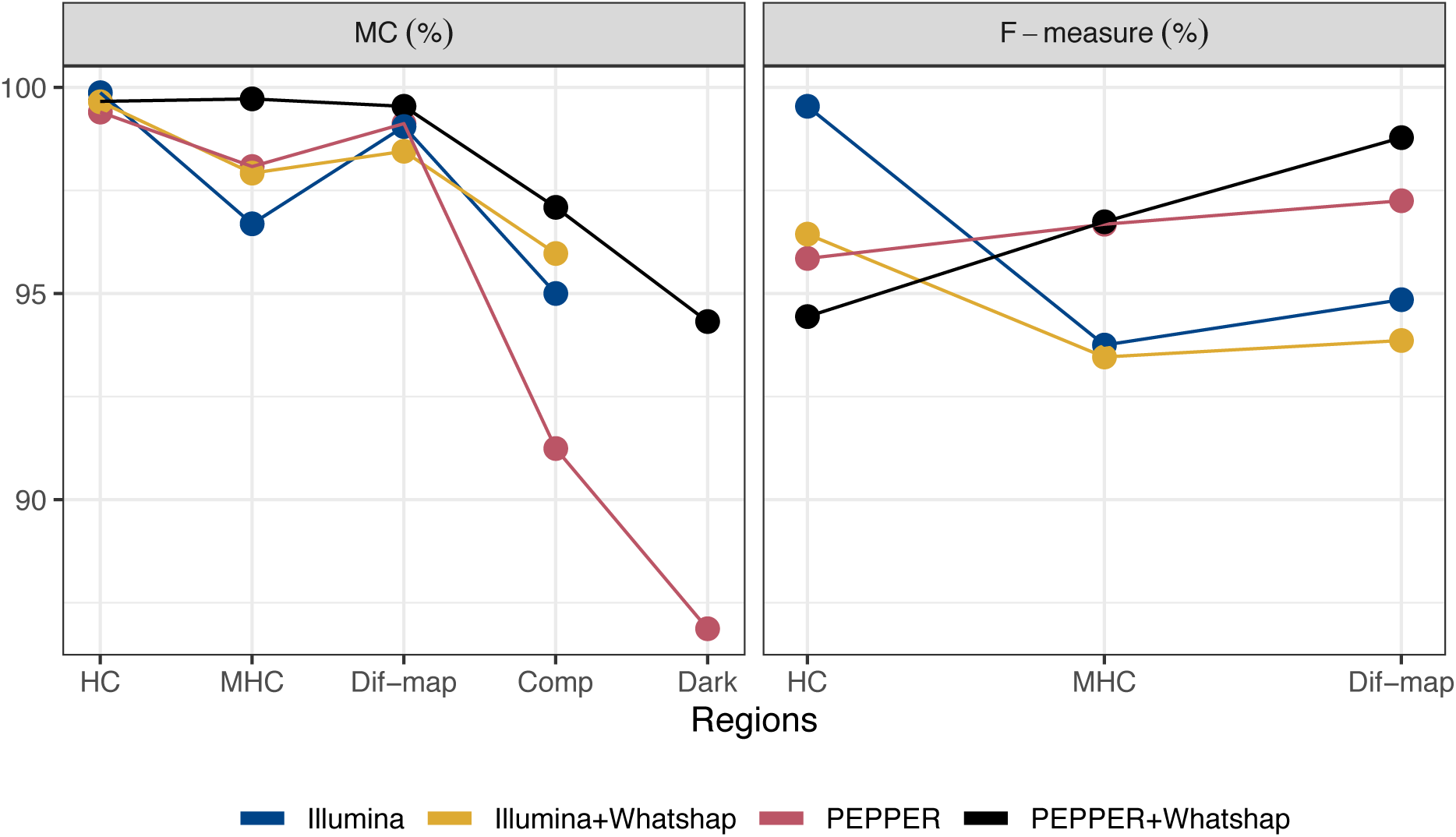
The effect of re-genotyping on mendelian concordance and F-measure. For PEPPER we see an increase in Mendelian concordance for all regions, while the F-measure is decreased in the HC, but increased in the MHC and difficult-to-map regions. For Illumina we see improved Mendelian concordance in the MHC and complementary regions, while it is decreased in the HC and difficult-to-map regions, the F-measure is decreased in all regions. HC: High confidence; MHC: Major histocompatibility complex; Dif-map: Difficult-to-map; Comp: Complementary regions (outside HC, excluding centromere and sex chromosomes).

For the Illumina data, the F-measure decreased for all regions (Figure 5), while re-genotyping had mixed effects on the Mendelian concordance with slight increases in the MHC and complementary regions and decreases in the HC and difficult-to-map regions.

## Discussion

We have analyzed multiple variant calling approaches in ONT data from the publicly available Ashkenazim trio (HG002, HG003, HG004). This has shown very consistent SNP calling, with both Medaka and PEPPER exceeding 99% F-measure when evaluating against the latest release of the GIAB truth sets. Limiting evaluations to more challenging regions of the truth sets shows performance 2-4% higher than SRS with as little as 20X coverage. No single variant calling approach was the best across all regions and coverages, but the consistency of PEPPER makes it a good default choice for analysis up to at least 50X coverage, while we begin to see diminishing results at 60X and above. Another benefit of PEPPER compared to other variant calling options is the ability to output gVCF format, which is easily processed with GLnexus [23,24] to create joint VCF files.

Evaluating performance outside the high confidence regions we observed decreased Mendelian concordance, which was expected as these regions are generally more challenging. Re-genotyping calls in these regions with Whatshap improved Mendelian concordance by almost 6%, resulting in ONT surpassing SRS. This approach also improved the F-measure of ONT data in both the MHC and the difficult-to-map regions. A similar approach for SRS was not beneficial, highlighting how some of the behavior of DeepVariant is good for short reads but at times detrimental to long reads. Re-genotyping PEPPER variant calls in the MHC (highest SNP-density region) also improved the consistency of variant calling, while maintaining a similar average F-measure, which in our opinion is to be preferred.

Finally, we show that ONT can reach an additional 22 mb of the genome, finding more than 50,000 variants, which are completely inaccessible to SRS. As shown, these regions include medically relevant genes like *CFC1B*, which can only be reached with the very long reads from ONT or potentially PacBio continuous long reads (CLR). During this work, we have observed new releases of almost every software used, including the Guppy basecaller used to generate the initial sequence files. It is therefore not unreasonable to expect improved performance of ONT data in the future, both through better basecalling but also variant calling. Meanwhile, SRS has reached a point where future improvements will be minimal, leading us to believe that the current performance difference in regions accessible to both methods will become smaller.

The primary issue for ONT is now indel performance, which we have not put substantial focus on in this paper. Current results are inferior to SRS in all evaluation regions by 5-30%, making it an obvious focus point for future development.

## Conclusion

For researchers whose primary interest is small variants inside the high confidence regions, SRS is both cheaper, better and easier to work with. ONT sequencing technology has already been shown to be useful for structural variant detection [25] and methylation calling [26]. We now show that ONT is beneficial for small variant calling in the MHC, the difficult-to-map regions and regions outside the high confidence regions, in particular we find 22 mb accessible only to ONT.

In the challenging regions of the genome, ONT outperforms SRS in SNP calling, helping researchers gain access to genomic regions and genes which are otherwise completely dark. Here we find PEPPER to be the best performing variant caller, without access to very high coverage data (>60X). Further, we advise a re-genotyping step, as it improves consistency and performance of variant calling in these regions.

As a technology, ONT is still quite immature, making it a challenge to utilize to its full potential. At the same time, this immaturity promises greater performance and easier use in the future, if developments continue at the current pace.

## Methods

### Data preparation

GRCh38 aligned Illumina BAM files from the Ashkenazim trio (HG002, HG003, HG004) were downloaded and used as is. ONT and PacBio HiFi FASTQ files were downloaded and aligned to GRCh38 using Minimap2 v2.14 [27] utilizing the presets for each technology (“-ax map-ont” and “-ax asm20”, respectively). Sorting and indexing was performed with SAMtools v1.9 [28].

All Illumina data had a coverage of 60X, ONT ranged from 50X (HG002) to 80X (HG003, HG004), while PacBio had 35X coverage for all.

### Variant calling

#### Illumina

A singularity image of DeepVariant v0.10.0 was created using ‘singularity pull deepvariant_0.10.0.simg docker://google/deepvariant:0.10.0’. Variant calling was performed with standard parameters (--model_type WGS) using the singularity exec command, outputting both VCF and gVCF files.

#### ONT

Medaka v1.0.3 was run using medaka_variant, calling variants by chromosome before combining VCF files with BCFtools v1.9 [28].

Clair v2.0.6 was run using the callVarBam module with the pretrained ONT model, variants were called by chromosome.

A singularity image of PEPPER/DeepVariant was created using ‘singularity pull docker://kishwars/pepper_deepvariant_cpu:latest’ (Image from 15/6/2020). Variants were called by modifying the run_pepper_deepvariant.sh script to include gVCF output and storing the PEPPER models outside the image to ensure write permission. Variant calling was performed in ‘Run-time’ mode.

For both data types gVCF output from each individual was joined using GLnexus v1.2.7. A singularity image was created using ‘singularity pull glnexus_v1.2.7.simg docker://quay.io/mlin/glnexus:v1.2.7’. The image was executed with --config DeepVariantWGS.

#### Re-genotyping

Whatshap v1.0 was used to re-genotype the joint VCFs from GLnexus, running Whatshap genotype on individual chromosome with the --indel flag. VCF files were re-combined using BCFtools concat. Whatshap genotype required the presence of a read group tag and sample in the bam header. This was added using SAMtools with the command ‘samtools addreplacerg -r “ID:HG002\tSM:HG002”’.

### Evaluation

#### Truth sets

Version 4.2 of truth sets (VCF + BED files) were downloaded for each individual and used with RTG tools v3.10.1 [29].

#### MHC and difficult-to-map regions

BED files of the major histocompatibility complex (MHC) and diffult-to-map regions, as defined by GIAB, were downloaded and used as is.

#### Outside high confidence regions

A complementary BED file of the HG002 high confidence regions was created using BEDTools v2.18.2 [30] to extract variants for mendelian concordance testing. From this BED file we subtracted the centromere regions (UCSC table browser > Mammal > Human > GRCh38 > All tables > centromeres) as well as the X and Y chromosome, as these regions were too noisy and unsuited for Mendelian concordance, respectively.

#### Dark regions and medically relevant genes

For Illumina, ONT and PacBio, dark regions were computed from the BAM files of the trio. Dark regions were defined as regions of at least 30 bp, with an average coverage below 5X or less than 10% of reads having a mapping quality at or above 10. The centromere regions were subtracted due to noise. Finally, for Illumina and PacBio we created BED files of dark regions reachable by ONT by subtracting the ONT BED file from each.

The overlaps between dark regions and genes were computed by intersecting the gene coordinates with the dark regions using BEDTools.

#### RTG Tools

To benchmark against the truth sets we evaluated each VCF file using the vcfeval module of RTG Tools, using the truth VCF as baseline (-b) and the truth BED to define the regions (--bed-regions). The evaluation region (-e) was defined using either the truth BED, the MHC BED or the difficult-to-map BED.

Benchmarking outside the truth sets was achieved by intersecting the joint VCF file from GLnexus with the “outside high confidence regions” BED, followed by RTG Tools mendelian to compute Mendelian concordance rates with both mother, father and mother+father. The same approach was used for other BED files when reporting Mendelian concordance for other regions.

#### Visualization

Visualizations were made with R v3.6 [31], ggplot2 [32], karyoploteR [33], inlmisc [34] and IGV v2.8 [35].

## Availability of data and materials

Illumina sequencing data and evaluation BED files are made available by the GIAB consortium [36–38] from their FTP server [39]. ONT and PacBio data are available through the FDA precision challenge [40]. Centromere BED file can be downloaded from UCSC [41,42] at [43]. The precomputed Clair model is available at [44]. Ensembl release 98 is available from their FTP server [45].

## Declarations

### Competing interests

G.H., D.B. and B.V.H. are employees of deCODE Genetics/Amgen, Inc.

### Authors’ contributions

P.L.M, G.H., D.B. and B.V.H. designed the experiments. P.L.M. performed all variant calling and benchmarking. G.H. developed the method for detection of dark regions. P.L.M wrote the initial version of the manuscript, and G.H., D.B., M.N. and B.V.H. contributed to subsequent versions. All authors reviewed and approved the final version.

## Acknowledgements

The authors would like to thank our colleagues from deCODE genetics and Amgen Inc. for their helpful feedback. We would also like to thank the individuals who provided a biological sample to Genome in a Bottle.

## Notes

### Competing Interest Statement

Guillaume Holley, Doruk Beyler and Bjarni V. Halldórsson are employees of deCODE genetics/Amgen Inc.

